# Pairwise Efficiency: A new mathematical approach to qPCR data analysis increases the precision of the classical calibration curve assay

**DOI:** 10.1101/399337

**Authors:** Yulia Panina, Arno Germond, Brit G. David, Tomonobu M. Watanabe, Laboratory for Comprehensive Bioimaging, RIKEN Center for Biosystems Dynamics Research(BDR)

## Abstract

The real-time quantitative polymerase chain reaction (qPCR) is routinely used for quantification of nucleic acids and is considered the gold standard in the field of relative nucleic acid measurements. The efficiency of the qPCR reaction is one of the most important parameters that needs to be determined, reported, and incorporated into data analysis in any qPCR experiment. The Minimum Information for Publication of Quantitative Real-Time PCR Experiments (MIQE) guidelines recognize the calibration curve as the method of choice for estimation of qPCR efficiency. The precision of this method has been reported to be between SD=0.007 (3 replicates) and SD=0.022 (no replicates). In this manuscript we present a novel approach to analysing qPCR data obtained by running a dilution series. Unlike previously developed methods, our method relies on a new formula that describes pairwise relationships between data points on separate amplification curves and thus operates extensive statistics (hundreds of estimations). The comparison of our method with classical calibration curve by Monte Carlo simulation shows that our approach can almost double the precision of efficiency and gene expression ratio estimations on the same dataset.

## INTRODUCTION

RT-qPCR is considered the most sensitive technique for nucleic acid quantification, enabling measurements on as little as several molecules of the target (1). The advantage of this method over earlier methods of quantification, such as end-point PCR followed by gel visualization, is the ability to account for the efficiency of the PCR reaction by following it in real time and gathering fluorescence data after each amplification cycle (2–4). The efficiency of the reaction is defined as the increase of product per cycle as a fraction of the amount present at the start of the cycle (5, 6), and it is assumed that the mean efficiency of a qPCR reaction is stable and maximal before reaction saturation. The reaction efficiency, due to the exponential nature of PCR, can have dramatic effects on quantification measurements. It has been estimated that an uncorrected 0.05 difference in amplification efficiency between a reference gene and a target gene can lead to a false estimation of the target gene expression change of 432% (7).

The calibration curve method is widely considered the most precise method for mean efficiency estimation (8) and is required by MIQE guidelines: “Calibration curves for each quantified target must be included with the submitted manuscript, slopes and y intercepts derived from these calibration curves must be included with the publication” (5). The calibration curve is built by creating a serial dilution of known DNA concentration and plotting the quantification cycle (Cq) values on the y-axis against the logarithm of the sample concentrations on the x-axis. The efficiency (E) is then estimated from the slope of this curve using the classical formula E=10^-1/slope^ - 1, and the estimation in this case is based on the knowledge of the concentrations of all diluted samples. However, due to insufficient precision of a single dilution set, it is usually recommended to run at least 3 PCR reaction replicates for each sample to have 3 dilution sets for a single calibration curve. It has been shown that replicating a calibration curve 3 times by this approach increases the precision of efficiency estimation expressed as a confidence interval (CI) from 8.3% to 2.3% (8). The downside of this approach is the increased workload and cost.

In attempts to overcome the increased workload problem, several other methods have been developed in the past decades to estimate qPCR efficiency and to improve qPCR precision in general, such as FPK-PCR (9), LinRegPCR (10), Cy0 (11) and others. However, according to a recent analysis by Rujiter and colleagues, the majority of these alternative methods are very similar in principle as they are based on determining the same basic parameters (called Fq, Cq and E) and “all calculate a target quantity using an efficiency value and a Cq value” (6). In addition, alternative methods that rely on different ways of approximating a single amplification curve have never yielded acceptable accuracy (12). Thus, it remains to be seen whether a truly novel approach could improve the precision of qPCR efficiency and ratio estimations.

Here, we present a mathematical approach that improves the precision of qPCR efficiency estimation, while reducing the necessary workload for qPCR. This approach does not rely on Cq values or amplification curve approximations. Instead, it defines pairwise relationships between fluorescence readings on several amplification curves of a dilution set. We employ three statistical steps to increase precision: 1) a new formula for E estimation, based on the relationship between data points on each of the amplification curves from the dilution series, 2) defining the boundaries of the curve region suitable for analysis, and 3) eliminating outliers in the process of data analysis. Because this approach is based on commonly used, robust statistical methods, it is systematic and can be applied in any setting and on any instrument as long as basic principles are conserved. Our results show that the application of this different mathematical approach makes it possible to almost double the precision in qPCR efficiency measurements without increasing the pipetting workload. In addition, we demonstrate a 2.3-fold improvement in precision of the gene expression ratio estimation. This constitutes a conceptual advance in the field of qPCR and allows for further development of ideas in this direction. Moreover, these advancements have important practical implications for those who use qPCR in their experimental work.

## MATERIALS AND METHODS

### DNA sample

Mouse embryonic stem cell line E14Tg2a was purchased from RIKEN Cell bank, JP (AES0135) and was maintained as previously described (13). Total RNA was extracted using RNeasy kit (Cat# 74106, Qiagen, Japan) following the manufacturer’s instructions. Genomic DNA digestion was performed on-column according to said instructions. RNA concentration and absorbance ratios (A_260/280_ and A_260/230_) were checked with a spectrophotometer Nanodrop 2000 Spectrophotometer (NanoDrop Technologies, Japan). To produce cDNA for qPCR analysis, 300 ng of total RNA were reverse-transcribed with an Omniscript RT Kit (Cat# 205111, Qiagen) in a total volume of 20 µl. The resulting DNA was assessed by spectrophotometric analysis and diluted to 100 ng/µl.

### Quantitative real-time PCR setup and reagents

qPCR was performed using a CFX96 Connect apparatus (Bio-Rad, Japan). Hard-Shell^®^ 96-Well PCR Plates (Cat # HSP 9601, Bio-Rad) sealed with optically clear adhesive seals (Microseal^®^ ‘B’ seal, Cat # MSB1001, Bio-Rad) were used in all experiments. The thermocycler program consisted of an initial hot start cycle at 95°C for 3 min, followed by 33 cycles at 95°C for 10 sec and 59°C for 30 sec. Mouse actin beta (Actb) was amplified using the following primers: F-5’- AACCCTAAGGCCAACCGTGAA-3’, R-5’-ATGGCGTGAGGGAGAGCATA-3’ (with estimated product length 194bp). The primers were used at a concentration of 300 nM. SYBR Green-based PCR supermix (Bio-Rad) was used for all reactions according to manufacturer’s instructions. Each reaction was performed in a final volume of 8 μL. To confirm product specificity, a melting curve analysis was performed after each amplification, and agarose gel analysis was performed to ensure the amplification of the right product (Supplementary Fig. S1).

### Experiment design and PCR dataset generation

For the assessment of precision of our method and comparison with the classical calibration curve method, we produced 16 replicas of a 6-step dilution series. We provide the detailed pipetting layout in Supplementary Fig. S2. Two datasets were generated from this experiment and processed using Bio-Rad CFX Manager 2.0 (2.0.885.0923). Dataset 1 consists of relative fluorescence data obtained from the aforementioned experiment: 6 serial dilution wells * 16 replicas = 96 wells. Fluorescence data in Dataset 1 are expressed as RFU (Relative Fluorescence Units) which is a term specific to Bio-Rad software. It is important to note that, since our goal was to improve the accuracy of the classical calibration curve, all RFU values were taken as already processed by Bio-Rad software with the same settings that were applied to the generation of Cq values, as follows: Baseline Setting set to Baseline Subtracted Curve Fit, Cq Determination Mode set to Single Threshold. Dataset 2 contains automatically generated Cq values corresponding to Dataset 1. The threshold was automatically set at 31.07 by the Bio-Rad software.

### Determination of the exponential region

The most suitable bounds of the exponential region of the respective amplification curves were determined experimentally (see Results). However, prior to the experimental estimation, we conducted an initial estimation using well-known conventional techniques, namely, the “first outlier” method, the First Derivative Maximum (FDM) and Second Derivative Maximum (SDM) approaches (9, 14)). Since the initial estimation was done solely in order to provide a general range for experimental testing, we chose the approaches mentioned above, even though other more sophisticated approaches have been suggested (9). The lower boundary of the exponential region has been defined as the point at which the signal significantly rises above the baseline level as determined by the formula of “first outlier” detection (14). The results of the formula application to the first calibration curve replica (wells A1 through A6) are provided in Supplementary Table S1. In agreement with these data, the tentative lower boundary of the exponential region was set at 10-40 RFU.

The FDM and SDM values for the first calibration curve replica can be found in Supplementary Table S2 and Supplementary Table S3. As expected, the values differ for samples with different initial DNA concentration, and are in the range of 17-23 cycles for SDM, and 18-25 cycles for FDM values. Supplementary Figure S3a shows the FDM values for the whole Dataset 1 plotted against cycle numbers. The earliest FDM was encountered at cycle 18 in the most concentrated sample. The latest FDM of the dataset came at cycle 25. As shown in Supplementary Figure S3b, the RFU values for cycles corresponding to calculated FDMs fall in the range of 120-230 RFU. Thus, in accordance with these data, the tentative initial estimation of the upper boundary of the exponential region to use in the experimental test was set between 120-230 RFU.

### Baseline treatment

Since the goal of our analysis was to directly improve the precision of the classical calibration curve method, the same software settings were applied to fluorescence data as to the generation of Cq values. The Bio-Rad software was set to Baseline Subtracted Curve Fit, and the baseline was subtracted automatically by the software producing Relative Fluorescence Unit values. This Bio-Rad subtraction method is based on either adding a constant value, or a linearly growing value to the raw fluorescence and thus does not eliminate the noise inherent to any qPCR instrument as an electric device.

### Evaluation of the noise influence

To determine the properties of noise and the scale of noise influence, we examined the fluorescence readings in the beginning cycles of the Dataset 1. As shown in Supplementary Fig. S4a, the fluorescence readings in the beginning cycles (up to cycle 13-18, depending on the starting concentration) were distributed close to 0, with inclusion of negative readings. The minimal value of the whole dataset was -9.44 RFU. To demonstrate the noise distribution, we show three histograms which contain fluorescence readings from the following cycles: 1) Cycles 1 through 5; 2) Cycles 1 through 10; and 3) Cycles 5 through 10. The data were taken from dataset 1 and two more 96-well plates replicating serial dilutions, with the Actb gene as target (raw data of these two plates are available on request). The total number of data points resulted in 2880 fluorescence readings (first 10 cycles from 96 wells in 3 plates). The result is shown in Supplementary Fig. S4b. The noise in the beginning cycles appeared to have a nearly normal distribution with a non-zero peak. The positions of the peaks and the distribution did not change depending on the number of included cycles, which indicated that there was no detectable signal at this stage - because the increasing signal would have produced a shift to the right in the noise distribution if it existed. Thus, we concluded that the initial fluorescence readings in our system contain noise, and the noise has the approximate range of -10 RFU to 10 RFU. To ensure that all data points that we would take for analysis contain the non-noise signal, we concluded that the lower boundary should not be lower than 10 RFU which is in accordance with the boundary set by the ‘first outlier’ (see Determination of the exponential region).

### Data processing

The data processing was carried out in Microsoft Excel and R. All excel files are available in supplementary materials.

## RESULTS

### Assessment of the stability of amplification efficiency in the exponential phase

The goal of our analysis was to increase the accuracy of measuring the mean amplification efficiency that is normally measured by the classical calibration curve method. According to the mainstream view, any PCR reaction proceeds with stable efficiency until end-stage reagent depletion and the accumulation of reaction products cause a steep decline in the efficiency, and the reaction gradually slows down (15, 16). The calibration curve method aims at measuring the mean stable efficiency of the reaction before the saturation occurs, and this mean maximal efficiency is assumed to be identical across all dilution samples. However, it has been argued that the sensitivity of modern PCR machines does not allow detection of a weak fluorescent signal in the exponential phase of the PCR reaction, where the efficiency is still stable, and the signal first appears when the efficiency is already declining (9, 17, 18). It has also been pointed out that the analyses based on mean efficiency should be conducted strictly at the region before efficiency decline, if such a region is detectable.

To determine if our system allows to detect the theoretical stable mean efficiency, we analysed the fluorescence readings data from Dataset 1 (see Materials and Methods for description) using the following formula for the calculation of mean efficiency E.

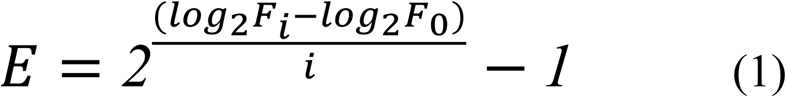

where *i* is the cycle number for a particular fluorescence reading F, and F_0_ is the initial fluorescence value of the sample. The logarithms, base 2, are used because the series contains 2- fold dilution sets.

The formula (1) cannot be used directly for E calculation because the fluorescence level of the starting material F_0_ is unknown. Thus, for the purpose of this analysis we have taken F_0_ values from the range of 0.007 to 0.0002 to substitute in the formula. This range was determined so that the resulting E roughly corresponded to the values obtained by the standard curve method (Supplementary Table S4). As shown in Fig. 1, we found that in the first cycles where the non-background signal is detected by the machine, E displays a relatively constant pattern (SD=0.01), while in the later cycles it starts to decline steadily (Supplementary Table S5). The initial region with the small standard deviation lasted from cycle 13 until cycle 17 for the most concentrated sample.

**Figure 1.**
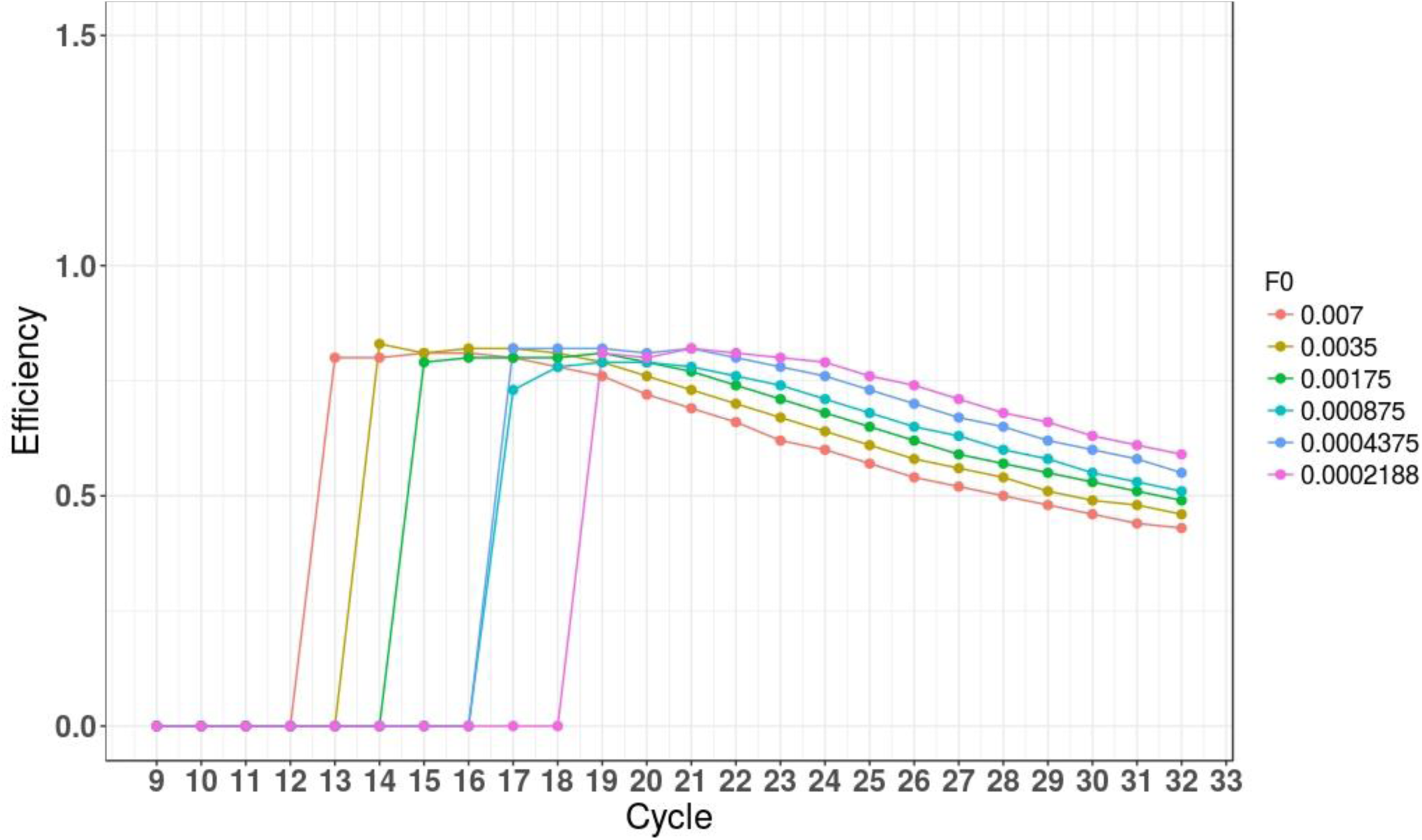
A graphical representation of mean efficiency (E) values across all cycles taken from a 6-step dilution set. Efficiency is calculated using the formula 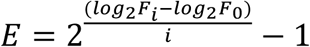. The F_i_ and i values for calculation are taken directly from Dataset 1, wells A1 through A6. Since F_0_ value is unknown, it was selected from the range of theoretically possible F_0_ values (covering 0.007 - 0.0002) and used in the formula.

Varying the F_0_ value did not affect the detection of this region of relatively constant E, as other curves also produced a similar pattern with small variation of E in the initial 4-5 cycles where the signal was already detected, and a steady decline after that.

According to these data, our experimental system allowed the detection of approximately 5 fluorescence values from the exponential phase of amplification where the variation of the mean efficiency does not exceed ±0.01. This result overall shows that the theoretical stable mean efficiency is detectable and can be quantified.

### Amplification efficiency estimation

Next, we approached the question of how to reduce the uncertainty in the mean E estimation given that only 5 or fewer fluorescence data points on each curve belong to the E stability region. For this purpose we introduced a new formula (4) for mean E estimation from a dilution set. This formula describes the relationship between 2 individual fluorescence readings in any given dilution set. The fluorescence readings are represented by data points on 6 amplification curves, in the case of one 6-step serial dilution experiment (Fig. 2b). The E estimation in our case is based on a relationship between a pair of actual fluorescence readings, as opposed to the slope of the calibration curve, which is based on cycle fraction values (Cq).

**Figure 2.**
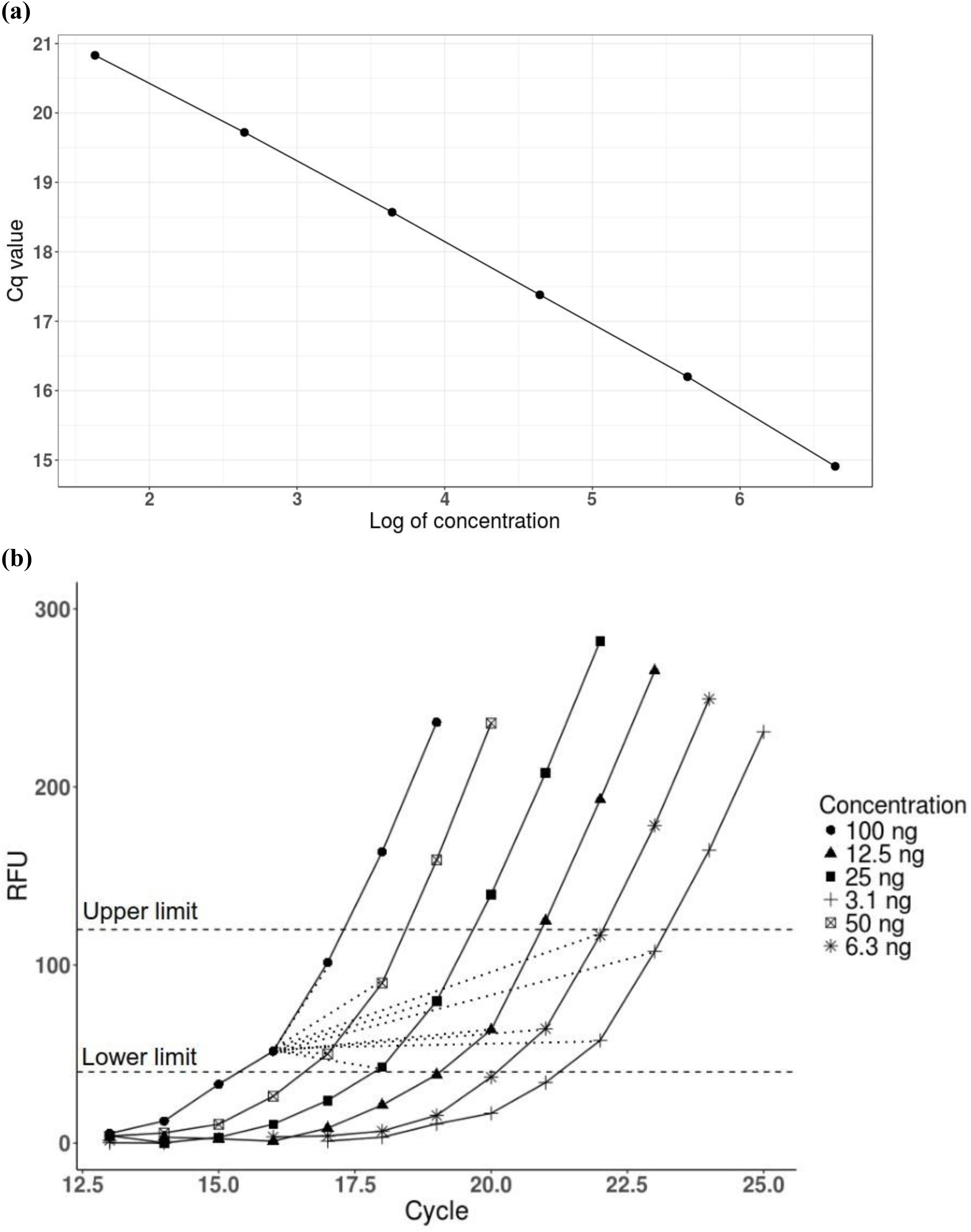
A graphical representation of the classical calibration curve method (a) and our new method (b). **(a)** The calibration curve was built from wells C1 through C6 in Dataset 2, which contained a serial dilution of the following concentrations: 100 ng - 50 ng - 25 ng - 12 ng - 6 ng - 3 ng. Logarithms of the concentration are plotted on the x-axis, and the corresponding quantification cycle (Cq) values are plotted on the y-axis. The lowest concentration corresponds to Cq=20.83, and the highest corresponds to Cq=14.91. The estimation of E was then performed by calculating the slope of the curve and applying the classical formula E=10^-1/slope^- 1, where E is dependent on the slope. **(b)** The amplification curves from the same wells C1 through C6 are shown (RFU data taken from Dataset 1). Different shapes (circles, squares, triangles etc.) represent fluorescence readings taken by the machine after each PCR cycle. The fluorescence increased with each cycle progressively. Cycle numbers are plotted on the x-axis, and Relative Fluorescence Units (RFU) are plotted on the y-axis. Horizontal lines denote the region of amplification curves from which the fluorescence data points were taken for analysis. In this example, the total of 11 fluorescence data points fall inside the denoted region. Efficiency estimation was then based on the pairwise analysis of the relationship of one data point to all other data points, as described by formula (4). Only one data point’s relationships are shown by straight lines reaching to other data points, for clarity.

When devising our formula, we used the same basic assumptions that the classical calibration curve method uses (6, 11) when calculating the mean efficiency on a calibration curve, namely:

1. The kinetics of a PCR reaction with a given DNA-primer set are the same irrespective of the initial template concentration.
2. The kinetics of the PCR reaction are assumed to be classical (described by the classical formula F=F0*(1+E)^i^)
3. The mean efficiency is maximal and constant before the reaction saturation.
4. Fluorescence readings and double-stranded DNA concentration are linearly related to each other, and the increase in fluorescence is directly proportional to the increase in target concentration.

Given these assumptions, any one fluorescence reading F on any one of the amplification curves in the dilution set can be described by the following equations:

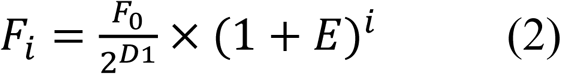

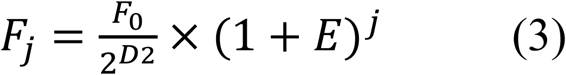

where i and j are cycle numbers for a particular fluorescence reading, F_i_ and F_j_are the fluorescence readings in cycle *i* or cycle *j*, F_0_is the initial fluorescence of the undiluted sample, D1 and D2 are dilution factors for curve 1 and curve 2 (if the pair of data points are on the same curve, then D1=D2), and E is the mean amplification efficiency for the qPCR reaction for the given DNA-primer set. The dilution factor D is defined as the logarithm of the fold-dilution, compared to the undiluted sample whose logarithm of the fold-dilution, by definition, is 0. Since we applied twofold dilutions for mathematical clarity, D values in our case were integers from 0 to 5. In the case of tenfold dilutions, the corresponding ‘2’ values in the formulae will become 10, and the dilution factors will remain unchanged.

The equations 2 and 3 allow us to calculate the mean efficiency for a given pair of fluorescence readings such as:

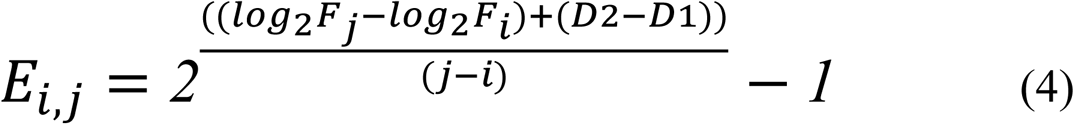

Thus, while the estimation of the mean efficiency across a dilution set by the calibration curve method is based on a single curve and produces a single E value, our new method calculates an array of E values based on all possible pair combinations from this dilution set, producing about 50-400 individual E measurements (depending on the number of fluorescence readings included in the exponential region taken for analysis), and then estimates the mean efficiency from this array of E measurements.

### Experimental determination of lower and higher boundaries

Because each dilution set produces amplification curves based on different concentrations of starting material, one can expect that the exponential region of each curve should start at a different cycle. Thus, it is necessary to experimentally determine the most suitable upper and lower boundaries of the exponential region for all curves taken together. An incorrect determination of the boundaries and subsequent inclusion of non-exponential values would be a major source of error in E estimation. To determine the most suitable boundaries, we experimentally tested at what fluorescence range (i.e. what portion of each of the amplification curves) the application of our method produces the results with the highest precision. For the estimation of precision we applied a Monte Carlo approach that has been previously described by Svec et.al. for the evaluation of precision of the calibration curve method (8). The lower boundary was tested at the range of 10 RFU - 80 RFU, and the higher boundary was tested at the range of 120 RFU - 230 RFU (see Materials and Methods). Using fluorescence readings from Dataset 1 that contained 16 technical replicas of a 6-step dilution set, we performed more than 100 samplings for each boundary combination, randomly selecting 1 out of 16 technical replicas for each concentration, allowing to test the majority of all possible combinations. Next, 50 separate mean E values were calculated based on random samplings of 3 values out of 16. This operation was repeated for various combinations of different boundaries, and the SD values of all mean E values obtained in each case were recorded (Fig. 3a). The exact SD values and other characteristics can be found in Supplementary Table S6.

**Figure 3.**
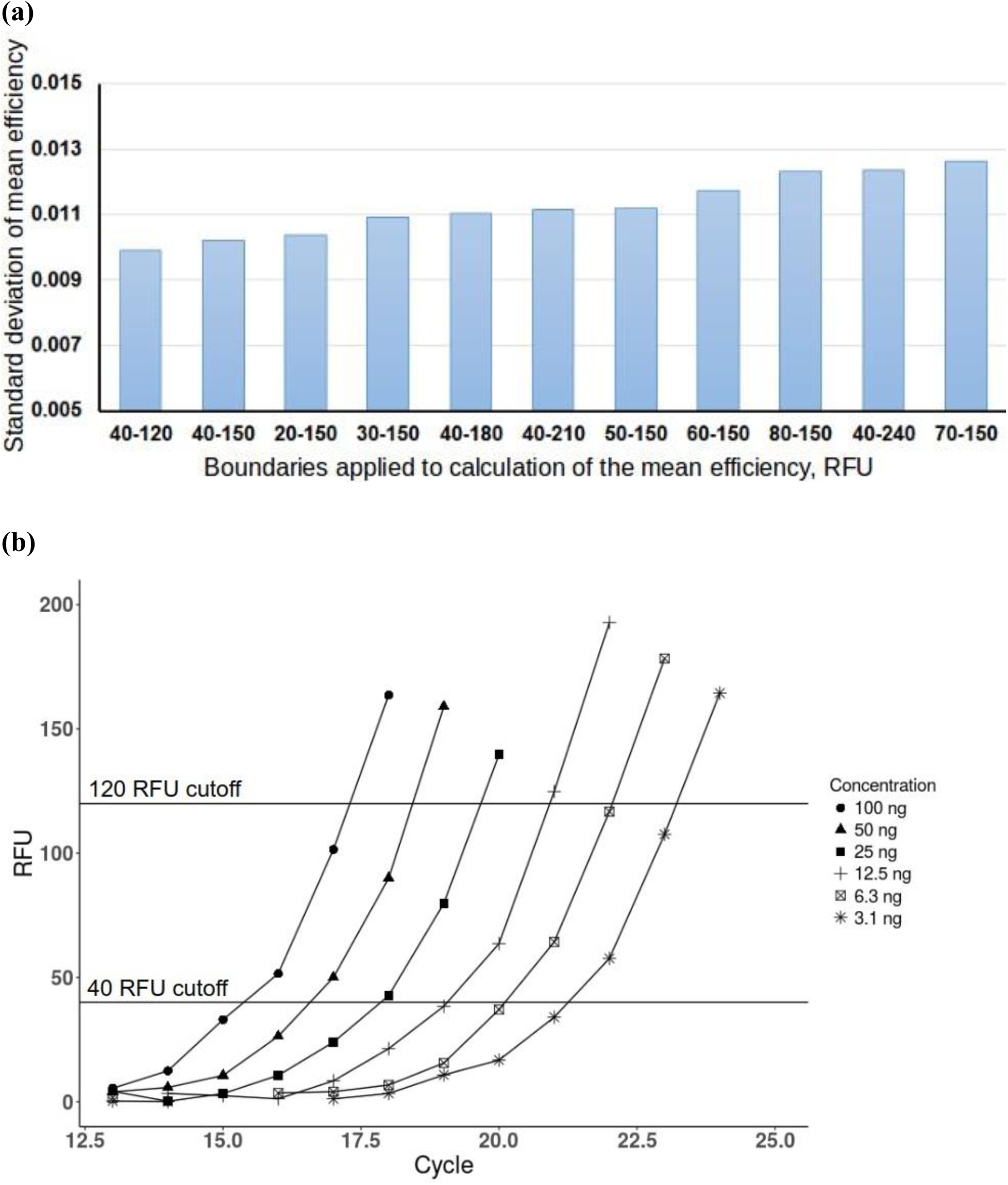
Determination of the most suitable RFU boundaries for a 6-step dilution series. **(a)** Standard deviations (SD) of the efficiency values calculated by a Monte Carlo approach using different regions of amplification curves. The mean efficiency (E) of a 6-step dilution set was calculated based on randomly selecting data out of 16 set replicas. Each time different portions of the amplification curves have been included in the calculations, defined by lower and upper boundaries. The lower boundary varied between 20 RFU and 80 RFU, while the upper boundary varied from 120 RFU to 240 RFU. The lowest SD was obtained when applying the following boundaries: lower at 40 RFU and upper at 120 RFU. The SD tended to rise when boundaries were raised. **(b)** The best-performing boundaries (40 RFU - 120 RFU) are depicted for one dilution set (in this case, wells A1 through A6). The best-performing boundaries are situated in the lower portion of amplification curves, just above the detection limit. 11 fluorescence readings fall inside the defined region.

While varying the boundaries within the exponential region (for definition of exponential region see Materials and Methods) did not produce a significant difference in SD values, the best result was, nevertheless, obtained at the lower portion of the curve (40-120 RFU) (Fig. 3a). The variation in the SD value did not exceed 0.001 for the lower portion (40-120 RFU, 40-150 RFU, 20-150 RFU). This result is in agreement with previous studies reporting that the threshold for measurements is best set at the lower portion of the amplification curve because the declining efficiency in later cycles might significantly affect the results (16). We conclude that slight variation of lower and upper boundaries in the range of about 30-150 RFU did not significantly affect the precision of E estimation. This is well in agreement with our data on exponential region estimation (see Materials and Methods) which puts the upper boundary at the fluorescence range 120-230 RFU and the lower boundary at the fluorescence range 20-40 RFU. (Supplementary Fig. S3 and S4).

### Statistical elimination of outliers

To further improve the precision of the mean E estimation, we applied the following strategy to eliminate the individual pairwise outlier E measurements. First, we tested the pairwise E measurements calculated for each dilution set for distribution normality in Dataset 1 (16 dilution set replicas in total). The skewness and kurtosis values were used to make the basic assessment of distribution normality. As shown in Table 1, the majority of skewness values significantly deviated from 0, signifying distribution asymmetry. In addition, all kurtosis values were positive, indicating that calculated pairwise E measurements from these dilution sets have a leptokurtic distribution. Thus, the parametric tools designed for normally distributed values, such as quartile ranges or sigma rules, cannot be applied in this case. Instead, we used the principles that are used for Pearson’s chi-squared test to exclude the outlier E values. According to the principles of the Pearson’s chi-squared test, when defining the goodness of fit in a dataset of more than 100 data points, those with a frequency of less than 5 should be excluded from analysis; i.e. the data points whose frequency is less than 5 are considered statistically unreliable. Thus, we have ensured that more than 100 individual pairwise E measurements are included in the analysis, and analysed each dilution set by these criteria. The pairwise E measurements with frequency less than 5 were considered outliers and were excluded from the calculation the mean E value of the dilution set.

**Table 1.** Estimation of distribution normality. The E values of 16 dilution sets were analysed for skewness and kurtosis. Skewness values that deviate from 0 indicate asymmetry of the distribution. Kurtosis values that deviate from 0 imply deviation from normal distribution. Since all kurtosis values calculated for these dilution sets are positive, they indicate that more data values concentrated near the mean value than would be expected in a normal distribution. The right column contains the numbers of individual pairwise E measurements for each dilution set that were taken for this analysis.

| Dilution set (wells) | Skew | Kurtosis | N of E measurements |
| --- | --- | --- | --- |
| A1-6 | 1.064 | 7.357 | 167 |
| B1-6 | 0.615 | 4.085 | 168 |
| C1-6 | 0.221 | 3.556 | 170 |
| D1-6 | 1.051 | 6.305 | 183 |
| E1-6 | 0.473 | 5.524 | 168 |
| F1-6 | 1.880 | 6.769 | 152 |
| G1-6 | 2.012 | 10.079 | 182 |
| H1-6 | 1.379 | 12.177 | 168 |
| A7-12 | -0.337 | 2.160 | 168 |
| B7-12 | 0.098 | 4.508 | 149 |
| C7-12 | 0.215 | 2.838 | 204 |
| D7-12 | 0.739 | 2.514 | 168 |
| E7-12 | 0.563 | 3.555 | 188 |
| F7-12 | -0.034 | 3.843 | 171 |
| G7-12 | 1.429 | 7.023 | 152 |
| H7-12 | -0.148 | 5.319 | 188 |

Thus, for example, the dilution set in wells A1 through A6 had 167 individual pairwise E measurements, skewness=1.06 and kurtosis=7.36. The frequency of E values below 5 was first encountered at E=0.6 (60% efficiency) on the lower end, and at E=1.05 (105% efficiency) on the higher end (Supplementary Table S6). Thus, all pairwise E measurements that exceeded 105% and did not reach 65% were excluded from the calculation of mean E for this dilution set. The mean E for wells A1 through A6 prior to outlier analysis was E=0.79, and after the removal of outliers became E=0.816. Other mean E values for the remaining 15 sets were processed on the basis of the same algorithm.

### Comparison of the new method with the calibration curve-based E estimation

Next, we set out to compare the precision of our method to the classical calibration curve method. The same Monte Carlo approach was used for comparison exactly as described in Svec et.al., with the important difference that our general population (Dataset 2) contained 16 replicas of the dilution series (96 wells) as opposed to 4 replicas (24 wells) described in the aforementioned paper. To evaluate the precision of E estimation by the classical calibration curve method, we took more than 1000 samplings, selecting 3 replicas for each concentration at random. Individual E estimations were made based on these randomly produced calibration curves, and standard deviation (SD) was calculated for the produced E values. The SD value for mean E estimation by the calibration curve was found to be 0.019. Next, we applied the same approach to the Dataset 1 that contains the corresponding RFU values, using randomly selected data points for E calculation by our method, as described in the previous section. The results are shown in Table 2. Our method allowed an increase in the precision from 0.010 to 0.019, thus nearly two-fold. While the average E values were found to be 80% in both methods, our method produced a smaller standard deviation and smaller difference between maximal and minimal E measurements. The dispersion of E values obtained by our method expressed as Max E - Min E did not exceed 0.045, as opposed to 0.072 obtained by the classical calibration curve. This means that the magnitude of random error in the E estimation was approximately two times lower in our method compared to the classical calibration curve.

**Table 2.** Comparison of the classical calibration curve method with the new method. The standard deviations (SD) obtained from the Monte Carlo test, maximal and minimal mean efficiency values, the range between maximal and minimal values, and the average efficiencies are shown. While the average mean E value was the same for both methods (E=0.80), the precision of the new method of E estimation, expressed as standard deviation (SD), was almost two times better, and the dispersion, expressed as the difference between maximal and minimal calculated E values, was 1.6 times smaller.

| Approach | SD | Max E | Min E | Max-Min difference | Average E |
| --- | --- | --- | --- | --- | --- |
| Calibration curve | 0.019 | 0.83 | 0.76 | 0.072 | 0.80 |
| New method | 0.010 | 0.82 | 0.78 | 0.047 | 0.80 |

Next, we investigated whether this increased precision in mean E estimation would translate into increased precision of gene expression ratio measurements. To do that, we calculated the magnitude of possible error for the classical calibration curve method and for our new method, using the same assumptions as described in Materials and Methods. For the calculation of expression ratios in the case of the classical calibration curve, we used the equations described by M. Pfaffl (19). The mathematical model presented in his publication is, in principle, equivalent to the model previously designed by Roche Diagnostics and takes into account the efficiency of both target and reference genes. The formula presented by Pfaffl has the following appearance:

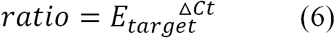

where ΔCt is the difference between Ct of the sample and Ct of control at the same threshold. Since our dataset of 16 dilution replicas contained exactly the same amount of target gene (Actb) in wells with the same concentration, theoretically the calculated ratio between these wells should be 1. Thus, we could measure the magnitude of the error in the determination of the ratio by measuring the maximal difference between each one of these 16 replicas. In this case, the error would be maximal when the efficiency value is maximal.

First, we determined which one of the 16 dilution sets gives the highest efficiency value. The analysis using the classical calibration curve method showed that wells D1 through D6 gave the highest efficiency (E=0.882). Next, using this efficiency, we applied the formula (6) for the undiluted samples, considering the Ct _sample_ the highest Ct from all 16 replicas, and Ct _control_ the lowest of all. This resulted in a ratio = 1.606. Thus, the maximal possible error in the estimation of a gene expression ratio when using the calibration curve method can reach up to 60%. Similarly, we used the maximal efficiency calculated by our method to estimate the magnitude of error on Dataset 1 with 16 replicas. The maximal efficiency value was obtained in the same wells (D1 through D6) as for the calibration curve, which underscores the robustness of both methods for E estimation. Using this maximal efficiency value, we estimated the F0 in all undiluted wells using our formula (2), based on actual fluorescence values. We obtained the following result: Max F=0.00435436, Min F=0.00345735. Then we calculated the difference between maximal F_0_ and minimal F_0_ which yielded a ratio=1.26. Thus, the magnitude of possible error in ratio estimation using our method amounts to 26%, which amounts to an improvement of about 2.3 fold in the precision of gene expression ratio estimation compared to the classical calibration curve method.

## DISCUSSION

qPCR is an affordable and widely used technique for nucleic acid quantification. However, despite its popularity, this method has yet to gain full trust in the biological community due to its limitations in providing precise measurements that may lead to low reproducibility. Multiple methods for qPCR data analysis have been developed throughout its history, yet the vast majority of them rely on Cq values, a calibration curve or curve fitting for efficiency estimation and subsequent data analysis, and do not achieve sufficient improvement in precision of efficiency or gene expression ratio estimations. Given this situation in the field, new approaches are needed that would allow a conceptual move forward. In this work, we provide a new approach to qPCR data analysis that consists of 3 elements. 1) It introduces a formula describing the relationship between two fluorescence readings on amplification curves, and does not rely on Cq values or a calibration curve for the mean efficiency estimation. 2) It estimates the boundaries of the exponential region for a group of amplification curves in order to determine reliable data boundaries. 3) It eliminates outliers in the process of calculating the mean E, as opposed to in the end.

qPCR is often associated with difficulties in reproducibility and excessive workload, such as creating multiple technical replicas to ensure statistical robustness. By contrast, our method provides a significant increase in the precision of efficiency and gene expression ratio estimation without increasing the workload. According to our analysis, 2 to 5 individual fluorescence readings from each amplification curve can be directly taken for the mean efficiency estimation. Six amplification curves from only six wells (which is 3 times less labor than required by calibration curve analysis) can provide 150-200 individual pairwise E measurements for far more extensive statistics. Thus, the workload necessary for achieving high precision can be substantially reduced.

Another advantage of our method is that it relies on actual fluorescence readings instead of implied data. It has been previously pointed out that the estimation of efficiency by the means of a classical calibration curve, as required by MIQE guidelines, is based not on the existent, but rather on implied data: “the data from a tube is discontinuous; fluorescence is measured at the end of each cycle, and there is no such thing as a fluorescence after a fractional number of cycles as implied by the continuous functions [that the classical Cq approach involves]”(20). We agree with this point of view. It can be said that one of the advantages of our method is that it is based on the analysis of actual fluorescence readings that are produced after each cycle, and does not rely on fractional cycles.

Finally, our method can be distinguished from other approaches because it allows the elimination of outlier values in the process of calculating the efficiency, and not at the end, as is done in other methods. For example, the MIQE guidelines require that the efficiency be estimated from the slope of the calibration curve, and considers efficiency value E to be the indicator of the robustness of the essay. In case if the E value exceeds the theoretical maximum of 100%, it is considered the result of reaction inhibition in one of the wells, and the whole assay usually needs to be repeated or redesigned (5). In contrast, because our method provides more than 150 individual pairwise E measurements for one replica of the calibration curve, it allows to apply both the distribution analyses for normality, and the application of the appropriate statistical instruments for outlier elimination. This aspect strongly differs from the more conventional methods where one or two “outlier” wells would often require the user to re-perform the whole experiment. In the case of our method, not only can we obtain more than 150 data points from a single dilution set (6 wells), but replication of the calibration curve 3 times would increase this number to 2556 (72 fluorescence readings, all in cross-pairwise relationships). This allows the use of powerful statistics, and it is a marked advantage of our method.

Overall, our method allows an almost two-fold increase in the precision of efficiency estimation and a 2.3-fold increase in precision of the gene ratio estimation (Table 2 and Results). Further developments of our approach, such as testing the application of different fitting and regression methods, can be explored. We hope that the provided approach will become a useful tool for the community and that our efforts will stimulate further investigations in improving the reliability of qPCR measurements.

## DATA AVAILABILITY

All data available in supplementary material.

